# Reinstatement of Pavlovian responses to alcohol cues by stress

**DOI:** 10.1101/2022.05.06.490952

**Authors:** Anne Armstrong, Hailey Rosenthal, Nakura Stout, Jocelyn M. Richard

**Affiliations:** Department of Psychological and Brain Sciences, Krieger School of Arts and Sciences, Johns Hopkins University, Baltimore, MD 21218; Department of Neuroscience, University of Minnesota, Minneapolis, MN 55415; Medical Discovery Team on Addiction, University of Minnesota, Minneapolis, MN 55415

**Keywords:** cues, alcohol, stress, addiction, rat, footshock, social defeat, yohimbine, Pavlovian conditioning

## Abstract

**Rationale:** Stress may contribute to relapse to alcohol use in part by enhancing reactivity to cues previously paired with alcohol. Yet, standard models of stress-induced reinstatement generally use contingent presentations of alcohol-paired cues to reinforce instrumental behaviors, making it difficult to isolate the ability of cues to invigorate alcohol-seeking.

**Objective:** Here we sought to test the impact of stress on behavioral responses to alcohol-paired cues, using a model of stress-induced reinstatement of Pavlovian conditioned approach, inspired by Nadia Chaudhri’s work on context-induced reinstatement.

**Methods:** Long Evans rats were trained to associate one auditory cue with delivery of alcohol or sucrose and an alternative auditory cue with no reward. Following extinction training, rats were exposed to a stressor prior to being re-exposed to the cues under extinction conditions. We assessed the effects of yohimbine, intermittent footshock and olfactory cues paired with social defeat on responses to alcohol-paired cues, and the effects of yohimbine on responses to sucrose-paired cues.

**Results:** The pharmacological stressor, yohimbine, enhanced Pavlovian responses to both alcohol and sucrose cues, but intermittent footshock and social defeat cues did not.

**Conclusions:** While yohimbine elicited reinstatement of Pavlovian conditioned responses, these effects may be unrelated to activation of stress systems.

## Introduction

Stress and alcohol-related cues contribute to alcohol use disorder in a variety of ways, including by triggering or exacerbating relapse of alcohol seeking and consumption (Brown et al. 1990; Lê et al. 1998; Sinha 2001; Logrip and Zorrilla 2012; Mantsch et al. 2015), and leading to escalated alcohol intake (Becker et al. 2011; Gomez et al. 2012; Hwa et al. 2015). Stress has been proposed to contribute to craving, consumption and relapse via many routes (Shaham et al. 2000; Sinha 2001), including by intensifying the motivational impact of drug-related cues (Robinson and Berridge 1993; Field and Powell 2007). Stress can also act as a contextual cue for the drug delivery or reward-seeking context. This stress context, like other reward-related environmental contexts (Zironi et al. 2006; Crombag et al. 2008), can then drive reinstatement of instrumental reward seeking (Schepers and Bouton 2019). Many studies of stress- or context-induced reinstatement of drug-seeking often involve response-contingent delivery of discrete drug-associated cues that are delivered throughout self-administration, extinction, and reinstatement testing (e.g. Tsiang and Janak 2006; Marinelli et al. 2007a; Bossert et al. 2007; Millan and McNally 2011). Because the same instrumental response was previously reinforced by simultaneous drug delivery and cue presentation, it is difficult in these studies to (1) determine the role of the cue in reinstatement per se and (2) to isolate the ability of cues to invigorate, as opposed to reinforce, reward-seeking actions.

A rich body of work led by Nadia Chaudhri and colleagues has shown that contextual cues can also renew responding for alcohol and other rewards in Pavlovian settings (Chaudhri et al. 2008b, 2013; Remedios et al. 2014; Sciascia et al. 2015; Valyear et al. 2017; Villaruel et al. 2017; Khoo et al. 2020; Segal et al. 2022). When rats undergo Pavlovian conditioning in one context and then have this Pavlovian association extinguished in a second context, placing rats back in the first context can drive reinstatement of responding to Pavlovian cues (Chaudhri et al. 2008b), similar to that seen for instrumental responding (Crombag et al. 2008; Chaudhri et al. 2008a; Janak and Chaudhri 2010). In addition to facilitating better understanding of the behavioral learning mechanisms that can lead to relapse-like behavior, this paradigm has facilitated neurobiological studies aimed at parsing the mechanisms underlying the invigoration of reward-seeking by environmental contexts, discrete cues, and interactions between these stimuli (Chaudhri et al. 2010, 2013; Valyear et al. 2020; Villaruel et al. 2022).

Inspired by these findings, and the possibility that stress potentiates alcohol use and relapse by potentiating the value of drug-paired cues, we sought to determine whether stress would similarly drive reinstatement of alcohol-seeking in a Pavlovian setting. We focused on stressors that have previously been shown to drive reinstatement of instrumental responding for alcohol. First, we assessed reinstatement of conditioned behavioral responses to alcohol and sucrose cues after exposure to a pharmacological stressor, yohimbine (Marinelli et al. 2007b; Lê et al. 2011; Bertholomey et al. 2016), an alpha-2 adrenoceptor antagonist. Next, we assessed the impact of a physical stressor, intermittent footshock, on behavioral responses to an extinguished alcohol cue (Lê et al. 1998). Finally, we evaluated the ability of a cue associated with a psychosocial stressor, social defeat in a resident-intruder paradigm (Funk et al. 2005; Manvich et al. 2015), to reinstate Pavlovian responding.

## Materials and Methods

### Subjects

Male and female Long Evans rats (n=43, 31 males and 13 females; Envigo) weighing 250-275 grams at arrival served as experimental subjects and were individually housed in a temperature-and humidity-controlled colony room on a 12 h light/dark cycle. A separate group of adult male Long Evans rats (n=4), weighing 500-750 at arrival served as resident aggressors in the social defeat experiment. These rats were each pair-housed with a sexually receptive, tubally-ligated adult female Long Evans rat (n=4 total) in a larger enclosure (approximately 3 × 2 ft) for nine days prior to social defeat conditioning. Rats trained with sucrose (n=8) were mildly food restricted to 90% of their free-feeding weight during training (∽5% of their body weight in chow was provided per day). All other rats were fed *ad libitum*, and water was provided *ad libitum* to all rats. All experimental procedures were approved by the Institutional Animal Care and Use Committees at Johns Hopkins University or the University of Minnesota and were carried out in accordance with the guidelines on animal care and use of the National Institutes of Health of the United States.

### Pavlovian conditioning with alcohol

Rats trained with alcohol reward (n=35) were first pre-exposed to ethanol in the homecage prior to training as described previously (Simms et al. 2008; Remedios et al. 2014; Millan et al. 2017). After one week of continuous access to 10% ethanol in the home cage, rats received chronic intermittent access to 20% ethanol in the homecage, for 24 hours at a time starting on Monday, Wednesday, and Friday, for a period of 7 weeks. All rats included here consumed at least 2g/kg/day of ethanol during the last two weeks of pre-exposure. Prior to cue conditioning, rats underwent port training in which they received 30 deliveries of .065 mL 20% ethanol to a reward port. Next, they underwent Pavlovian conditioning (20-25 sessions; MWF) as described previously (Ottenheimer et al. 2019). Rats were randomly assigned one of the following cues as their conditioned stimulus (CS+) for training and testing: white noise or a 2900 Hz tone. Rats received the alternate auditory cue as their CS-. During conditioning sessions, the CS+ and CS-, each lasting 10s, were presented on a pseudorandom variable interval schedule with a mean intertrial interval (ITI) of 80s. At 9s after the CS+ onset, 0.065 mL of 20% ethanol was delivered into the reward delivery port over a period of 1s. Rats then underwent extinction training and reinstatement testing with either yohimbine (n=17), intermittent footshock (n=11) or social defeat cues (n=7).

### Pavlovian conditioning with sucrose

Prior to cue conditioning, rats trained with sucrose reward (n=8 males) underwent port training in which they received 30 deliveries of .13 mL 10% sucrose to a reward port. They then underwent 10 sessions of Pavlovian conditioning with sucrose as described previously (Richard et al. 2018; Ottenheimer et al. 2019). During sucrose conditioning sessions the CS+ and CS-, each lasting 10s, were presented on a pseudorandom variable interval schedule with a mean intertrial interval (ITI) of 45s. At 8s after the CS+ onset, 0.13 mL of 10% sucrose was delivered into the reward delivery port over a period of 2s. Following Pavlovian conditioning with sucrose rats underwent extinction and reinstatement testing with yohimbine.

### Reinstatement testing with the yohimbine

Following conditioning with sucrose (n=8 males) or ethanol (n=17; 10 males and 7 females), rats underwent 5 extinction sessions in which cues were presented, but sucrose or ethanol reward was no longer delivered. Next, rats underwent a reinstatement session in which they either received an injection of yohimbine (2 mg/kg) or injection of saline vehicle, each given at a volume of 1ml/kg. Rats were placed in the operant chambers for 30 minutes prior to being re-exposed to 30 CS+ and CS- presentations under extinction conditions. Most rats (all sucrose, 10/17 alcohol rats) received a second reinstatement session 3-5 days later in which they received the opposite injection 30 minutes prior to being presented with 30 CS+ and CS-cues under extinction conditions.

### Conditioned reinforcement testing with yohimbine

To assess whether yohimbine enhances the value of Pavlovian-conditioned sucrose cues, rats that were previously tested for yohimbine reinstatement with sucrose cues underwent 3 additional days of Pavlovian conditioning prior to a conditioned reinforcement test. Rats received injections of yohimbine (2 mg/kg) or injection of saline vehicle, each given at a volume of 1ml/kg. Following 30 minutes in their home cages, rats were placed in the operant chambers for the conditioned reinforcement test, which lasted 45 minutes. During each session, entries into one nosepoke port resulted in 2s presentations of the CS+ on a fixed-ratio-2 (FR2) schedule. Entries into the other nosepoke port resulted in 2s presentations of the CS-on an FR2 schedule. Nosepokes during cue presentations were recorded but had no programmed consequences. Rats underwent a second session with the opposite injection, counterbalanced for order.

### Reinstatement testing with intermittent footshock

Following Pavlovian conditioning with alcohol, rats (n=11; 6 males and 5 females) underwent 5 extinction sessions in which cues were presented but alcohol was no longer delivered. On the reinstatement test day rats received 10 minutes of intermittent footshock (ITI ∽= 45 sec) (Le et al. 1999) or waited for 10 minutes in the operant chambers. Rats were assigned either 0.4 mA (n=5) or 0.8 mA (n=6) footshock intensity, which were tested on a between-subjects basis. At the end of the footshock or waiting period, rats were re-exposed to 30 CS+ and CS-presentations under extinction conditions. Rats were tested again 3-5 days later under the conditions they had not previously received (footshock versus waiting).

### Social defeat conditioning and reinstatement testing

Following Pavlovian conditioning with ethanol, male rats (n=7) underwent four social defeat conditioning sessions using the resident-intruder paradigm as described previously (Funk et al. 2005; Manvich et al. 2015). Pavlovian conditioned subjects (intruders) were placed inside a resident male aggressor’s home cage from which the female rat had been temporarily removed. Prior to the interaction the cage was prepared with either a lemon or banana odor on a cotton ball, counterbalanced across subjects. The resident rats’ home cages (approximately 3×2 feet) were constructed so that a perforated Plexiglas barrier could be inserted to bisect the box into two sides from which the rats could see, smell, and hear each other, but had limited physically interaction. The interaction was terminated, and the barrier inserted between the two rats once (1) the intruder rat sustained 3 bites from the territorial rat, (2) the intruder engaged in a supine submissive posture for 4s, or (3) 4 minutes had elapsed. After the barrier was inserted between the rats, the rats stayed in the behavioral box for at least one more minute, or until 5 total minutes had elapsed. Intruder rats faced a different resident rat for each interaction but were exposed to the same scent for each social defeat session. After each social defeat session, the intruder rats spent five minutes alone in a separate behavioral box with a cotton ball saturated with the scent that was not their respective social defeat scent.

The social defeat training was followed by five days of extinction training, during which Pavlovian conditioned rats were placed in the operant chambers under conditions identical to training, but the cues no longer predicted the delivery of alcohol, or lack thereof. Next, rats underwent reinstatement tests. During each test rats were either exposed to their social defeat scent or the control scent via a scented cotton ball that was placed in the operant chamber. They were then presented with 30 presentations each of the CS+ and CS-cues. In the subsequent session rats were tested with the opposite scent. Only one scent (lemon or banana) was tested in the operant chambers each day.

## Results

### Yohimbine reinstatement of responses to alcohol cues

Across 20-25 training sessions, rats learned to discriminate between a CS+ paired with 20% ethanol delivery and an unpaired CS-cue. Port entries during the CS+, but not CS-, increased across training days (Figure 1A; main effect of session, F(1,594) = 85.616, p < 0.001; interaction of session and cue type, F(1,594) = 49.098, p < 0.001). Port entries during the intertrial interval decreased across training sessions (main effect of session, F(1,297)=9.092, p = 0.0028). Rats then underwent 5 extinction sessions in which ethanol was no longer delivered. Across these sessions port entries decreased during both cue types (Figure 1B; main effect of session, F(1,136)= 39.735, p < 0.001), but more so during the CS+ (interaction of session and cue type, F(1,136) = 7.716, p = 0.0062). Despite the greater decrease in CS+ port entries, rats continued to discriminate between these cues, even during the final extinction session (CS+ versus CS-, F(1,10)=7.36, p = 0.022). Port entries during the intertrial interval also decreased across extinction training (main effect of session, F(1,69)=10.79, p = 0.0016).

**Figure 1.**
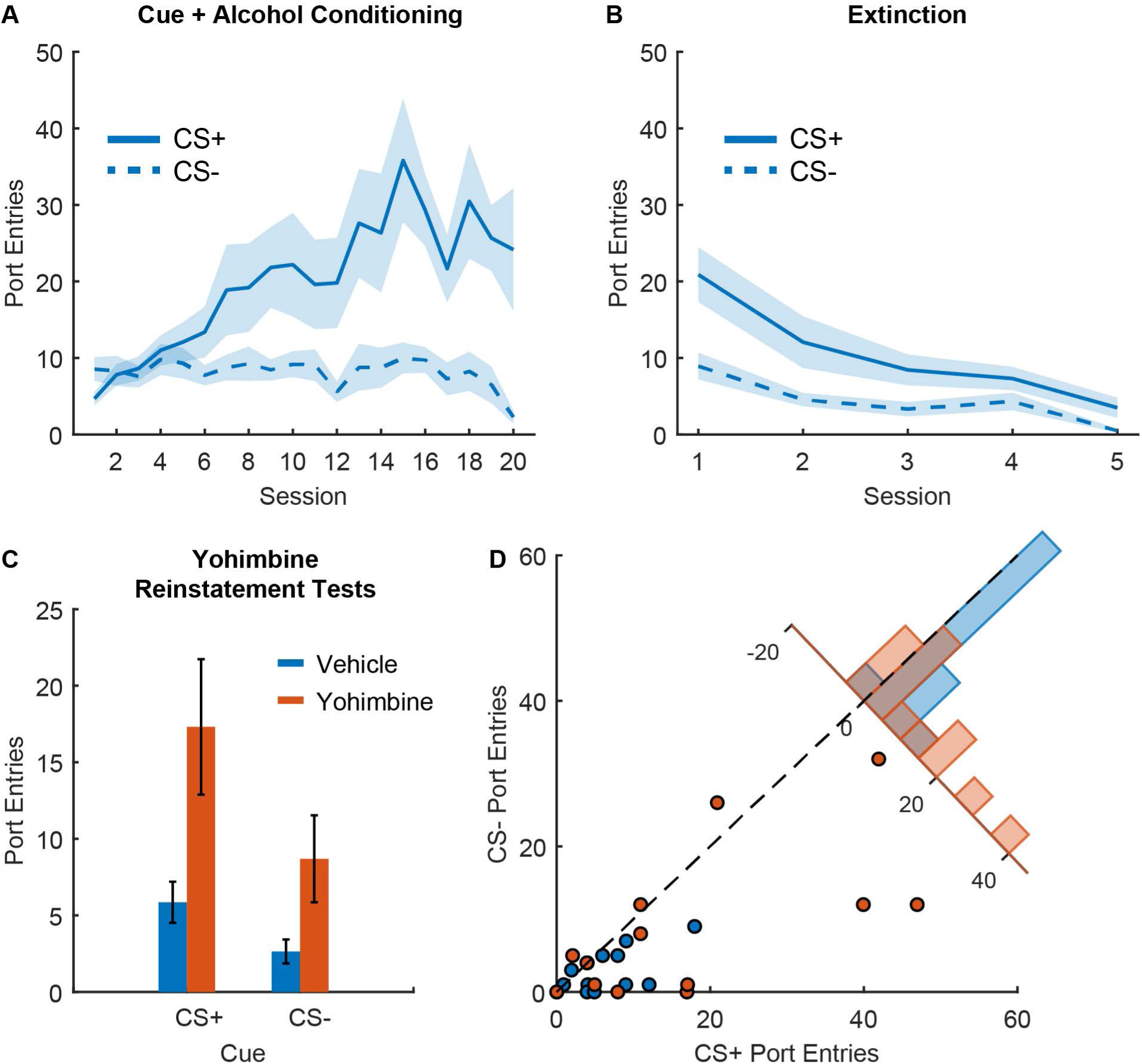
Yohimbine reinstatement of responses to alcohol cues. A) Port entry number during the CS+ and CS-across Pavlovian conditioning with 20% ethanol reward (n=17), mean +/- SEM. B) Port entry number during the CS+ and CS-across extinction training, mean +/- SEM. C) Port entry number during reinstatement tests during the CS+ and CS- after injection of yohimbine (orange) or vehicle (blue), mean +/- SEM. D) Scatterplot showing port entry number during the CS+ and CS-from individual reinstatement sessions after injections of yohimbine (orange) or vehicle. Inset histogram shows the distribution of sessions based on the difference in the number of port entries during the CS+ and the CS-.

We next assessed the ability of yohimbine to reinstate port entry behavior during ethanol cues. In comparison to a saline control session, we found that yohimbine significantly enhanced port entries during the cues (Figure 1C; main effect of yohimbine, F(1,50)=14.429, p < 0.001), but that this response was not selective to the CS+ (interaction between yohimbine and cue type, F(1,50)=2.156, p = 0.148), and we did not observe a significant effect of cue type (F(1,50)=1.59, p = 0.214), though the difference between CS+ and CS-responses was significantly different from zero (Figure 1D; F(1,25)=4.25, p = 0.049). We also assessed the impact of yohimbine on the difference between port entries during the CS+ versus CS-; though we observed a numerical increase following yohimbine, this comparison did not reach statistical significance (F(1,25) = 2.598, p = 0.119). Additionally, we also observed a significant effect of yohimbine on port entries during the intertrial interval (F(1,25)=6.698, p = 0.0158), suggesting that yohimbine increased port entries across the session, and not selectively during the CS+.

### Yohimbine reinstatement of responses to sucrose cues

We next sought to determine whether yohimbine would also reinstate port entries after Pavlovian conditioning with 10% liquid sucrose. Across 10 sessions of training, rats learned to discriminate between a CS+ paired with sucrose delivery, and an unpaired CS- (Figure 2A; main effect of session, F(1,156)=25.538, p < 0.001; interaction of session and cue type, F(1,156)=12.275, p < 0.001). Rats then underwent 5 extinction sessions in which sucrose was no longer delivered (Figure 2B). Across these sessions port entries decreased during the CS+ but not the CS- (main effect of session, F(1,76)=44.50, p < 0.001; interaction of session and cue type, F(1,76)=22.914, p < 0.001). By the final day of extinction rats no longer demonstrated significant discrimination between the CS+ and CS- (F(1,14) = 3.274, p = 0.092).

**Figure 2.**
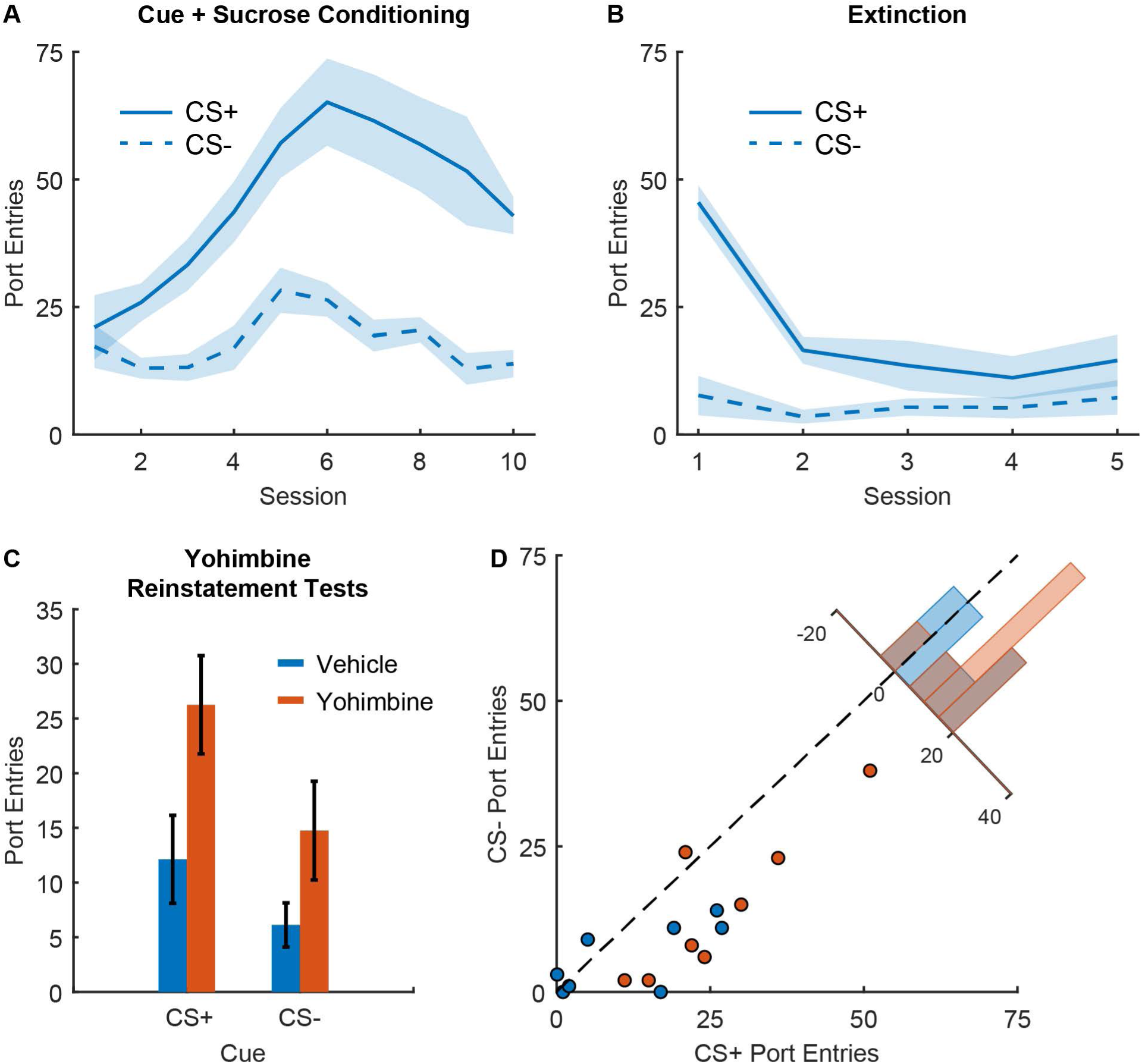
Yohimbine reinstatement of responses to sucrose cues. A) Port entry number during the CS+ and CS-across Pavlovian conditioning with 10% sucrose reward (n=8), mean +/- SEM. B) Port entry number during the CS+ and CS-across extinction training, mean +/- SEM. C) Port entry number during reinstatement tests during the CS+ and CS- after injection of yohimbine (orange) or vehicle (blue), mean +/- SEM. D) Scatterplot showing port entry number during the CS+ and CS-from individual reinstatement sessions after injections of yohimbine (orange) or vehicle. Inset histogram shows the distribution of sessions based on the difference in the number of port entries during the CS+ and the CS-.

We next assessed the ability of yohimbine to reinstate port entry behavior during sucrose cues. During the reinstatement tests, we found that yohimbine significantly enhanced port entries during the cues (Figure 2C; main effect of yohimbine, F(1,28)=23.916, p < 0.001) though this effect was not selective to the CS+. Specifically, we found a main effect of cue type (F(1,28) = 4.32, p = 0.047), but no interaction with treatment condition (F(1,28) = 1.813, p = 0.189). We also assessed the impact of yohimbine on the difference between port entries during the CS+ versus CS-; though we observed a numerical increase following yohimbine, this comparison did not reach statistical significance (Figure 2D; F(1,14) = 2.49, p = 0.137). Additionally, we also observed a significant effect of yohimbine on port entries during the intertrial interval (F(1,14) = 23.48, p < 0.001), suggesting that yohimbine increased port entries across the session, and not selectively during the CS+.

### Yohimbine effects on conditioned reinforcement by a sucrose cue

While yohimbine-induced increases in port entry behavior were not specific to the CS+ phase for rats trained with sucrose or alcohol, the cues may have still impacted port entry behavior during the session more generally. Therefore, we next sought to determine whether yohimbine would impact other aspects of cue-related behavior by assessing conditioned reinforcement. Yohimbine has previously been shown to increase conditioned reinforcement by cues predicting both 12% ethanol and 21.7% sucrose (Tabbara et al. 2020). Rats tested for yohimbine-induced reinstatement of Pavlovian sucrose-seeking were retrained with the CS+ paired with sucrose, and then underwent conditioned reinforcement testing. Overall we found that rats made more entries into a nose poke that produced CS+ presentations in comparison to a nose poke that produced CS-presentation (Figure 3A; main effect of cue type, F(1,28) = 8.207, p = 0.0078), but that yohimbine did not significantly impact this behavior (Figure 3C; main effect of yohimbine, F(1,28) = 2.447, p = 0.128; interaction between yohimbine and cue type, F(1,28) = 10.0649, p = 0.31). While we observed a numerical increase in port entries during the yohimbine session, this effect was not significant (Figure 3B; F(1,14) = 3.6929, p = 0.075). We also observed no relationship between CS+ nose pokes and total port entries (Figure 3B; F(1,12) = 0.879, p = 0.367) or any effect of yohimbine on this relationship (F(1,12) = 0.204, p = 0.659). Overall yohimbine did not produce a reliable increase in conditioned reinforcement.

**Figure 3.**
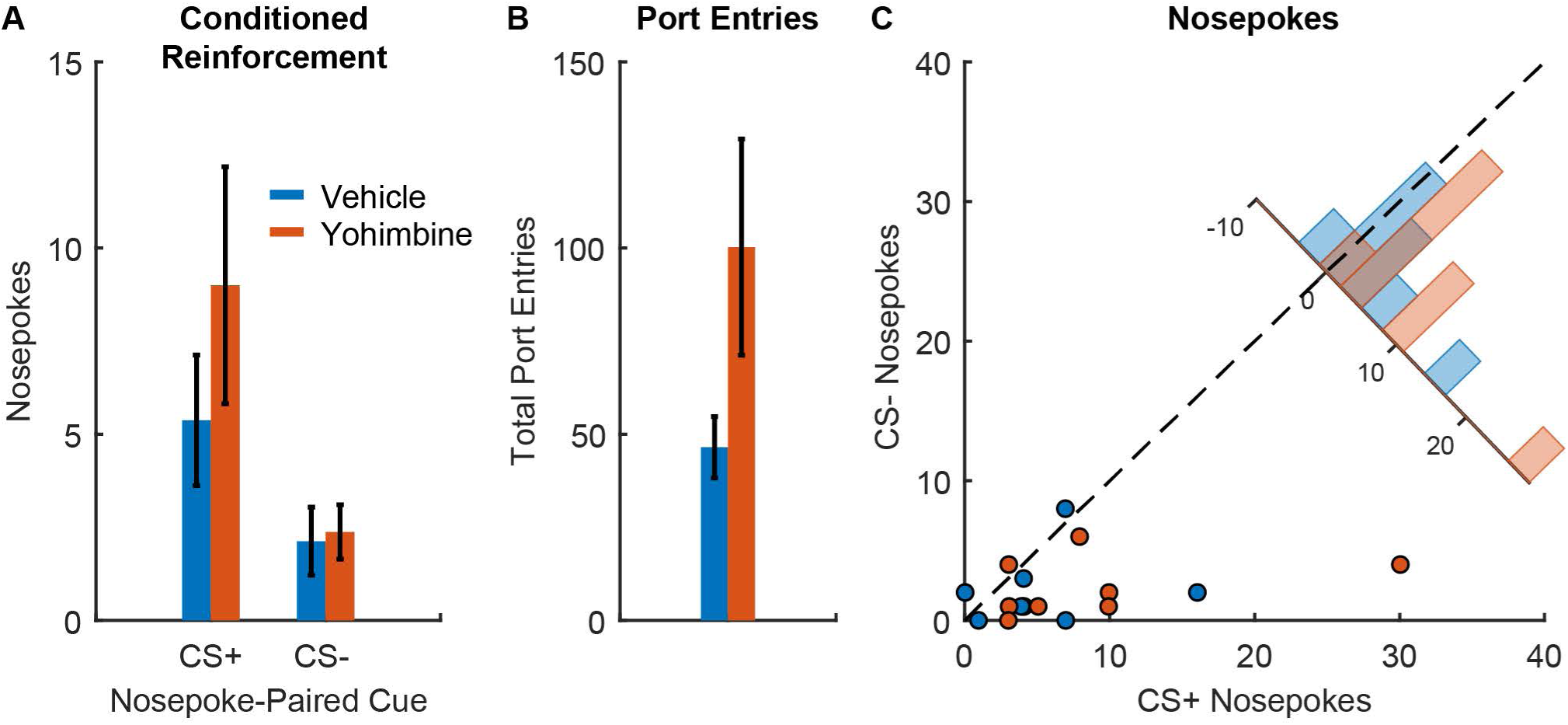
Effects of yohimbine on conditioned reinforcement by sucrose cues. A) Number of entries into nosepoke holes that resulted in presentations of the CS+ versus CS-following injections of yohimbine (orange) versus vehicle (blue) in rats that underwent Pavlovian conditioned with 10% sucrose reward (n=8), mean +/- SEM. B) Total port entries during conditioned reinforcement sessions after yohimbine (orange) versus vehicle (blue), mean +/- SEM. C) Scatterplot showing entries into the CS+-paired nosepoke versus the CS- nosepoke from individual sessions after yohimbine (orange) or vehicle (blue). Inset histogram shows the distribution of sessions based on the difference in the number of CS+ versus CS-nosepokes.

### Footshock reinstatement of responses to alcohol cues

We next sought to determine the impact of a physical stressor, intermittent footshock, on behavioral responses to an extinguished alcohol CS+. Rats first learned to discriminate between a CS+ predicting ethanol and an unpaired CS- (Figure 4A; main effect of session, F(1,454) = 81.783, p < 0.001; main effect of cue type, F(1,454) = 0.10924, p = 0.74; interaction of session and cue type F(1,434)=44.989, p < 0.001). Rats then underwent extinction sessions in which ethanol was no longer delivered, and port entry behavior then decreased (Figure 4B; main effect of session, F(1,106) = 36.746, p < 0.001, main effect of cue type, F(1,106)=51.548, p<0.001; interaction of session and cue type, F(1,106) = 14.889, p < 0.001). While port entry behavior decreased selectively during the CS+ relative to the CS-, rats continued to discriminate between these cues even on the final day of extinction (F(1,20) = 10.493, p = 0.0041).

**Figure 4.**
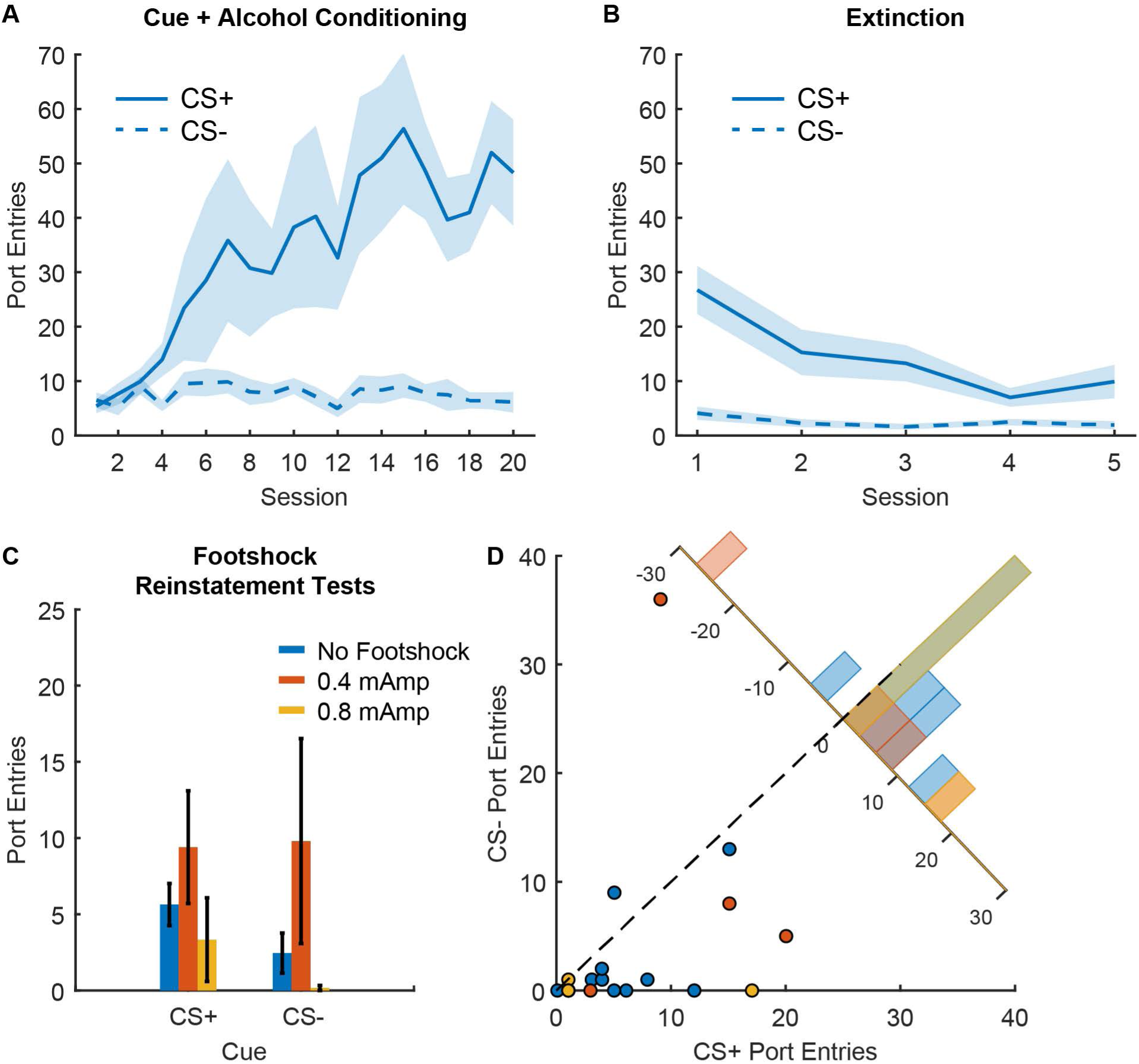
Footshock reinstatement of responses to alcohol cues. A) Port entry number during the CS+ and CS-across Pavlovian conditioning with 20% ethanol reward (n=11), mean +/- SEM. B) Port entry number during the CS+ and CS-across extinction training, mean +/- SEM. C) Port entry number during reinstatement tests during the CS+ and CS-after 0.4 mAmp intermittent footshock (orange), 0.8 mAmp intermittent footshock (yellow) or a 10-min control period without footshock (blue), mean +/- SEM. D) Scatterplot showing port entry number during the CS+ and CS-from individual reinstatement sessions after 0.4 mAmp intermittent footshock (orange), 0.8 mAmp intermittent footshock (yellow) or a 10-min control period without footshock (blue). Inset histogram shows the distribution of sessions based on the difference in the number of port entries during the CS+ and the CS-.

We then assessed the impact of intermittent footshock on subsequent port entry behavior during the alcohol cues (Figure 4C). In contrast to yohimbine, we found no effect of footshock on port entries during the cues (main effect of footshock, F(2,38) = 0.898, p = 0.415) regardless of cue type (interaction of footshock and cue type, F(2,38) = 0.342, p = 0.71). We also found no evidence of cue discrimination during the test (main effect of cue type, F(1,38) = 1.54, p = 0.222) or effect of footshock on cue discrimination (Figure 4D; F(2,19) = 0.473, p = 0.63). While we observed a numerical increase in total port entries after the 0.4 mAmp footshocks, this effect did not reach statistical significance (F(2,19) = 2.80, p = 0.085). Overall, intermittent footshock stress did not produce reliable increases in alcohol-seeking behavior, including in response to cues.

### Impact of a social defeat cue on behavioral responses to alcohol cues

Next, we sought to determine the impact of a psychosocial stressor on behavioral responses to alcohol cues. Because acute social defeat has been shown to reduce alcohol self-administration and responding in a reinstatement test (Funk et al. 2005), we elected to use odor cues paired with social defeat, which have been shown to produce reinstatement of instrumental responding for both alcohol and cocaine seeking (Funk et al. 2005; Manvich et al. 2015). Rats first learned to discriminate between a CS+ auditory cue predicting alcohol delivery and an unpaired CS- (Figure 5A; main effect of session, F(1,276) = 63.425, p < 0.001; main effect of cue, F(1,276)=0.142, p = 0.706; interaction of session and cue type, F(1,276) = 28.517, p < 0.001). Following the final 4 sessions of Pavlovian conditioning, rats underwent 4 social defeat sessions paired with their assigned odor cue. Next, rats underwent extinction training, during which their port entry behavior decreased across sessions (Figure 5B; main effect of session, F(1,108) = 17.976 p < 0.001), moreso during the CS+ than the CS- (main effect of cue type, F(1,108) = 30.437, p < 0.001; interaction of session and cue type, F(1,108) = 9.062, p = 0.0032). While CS+ port entries decreased more than CS-port entries, port entries during the CS+ remained numerically higher on the final day of extinction, though this difference was not statistically significant (F(1,12) = 4.18, p = 0.063).

**Figure 5.**
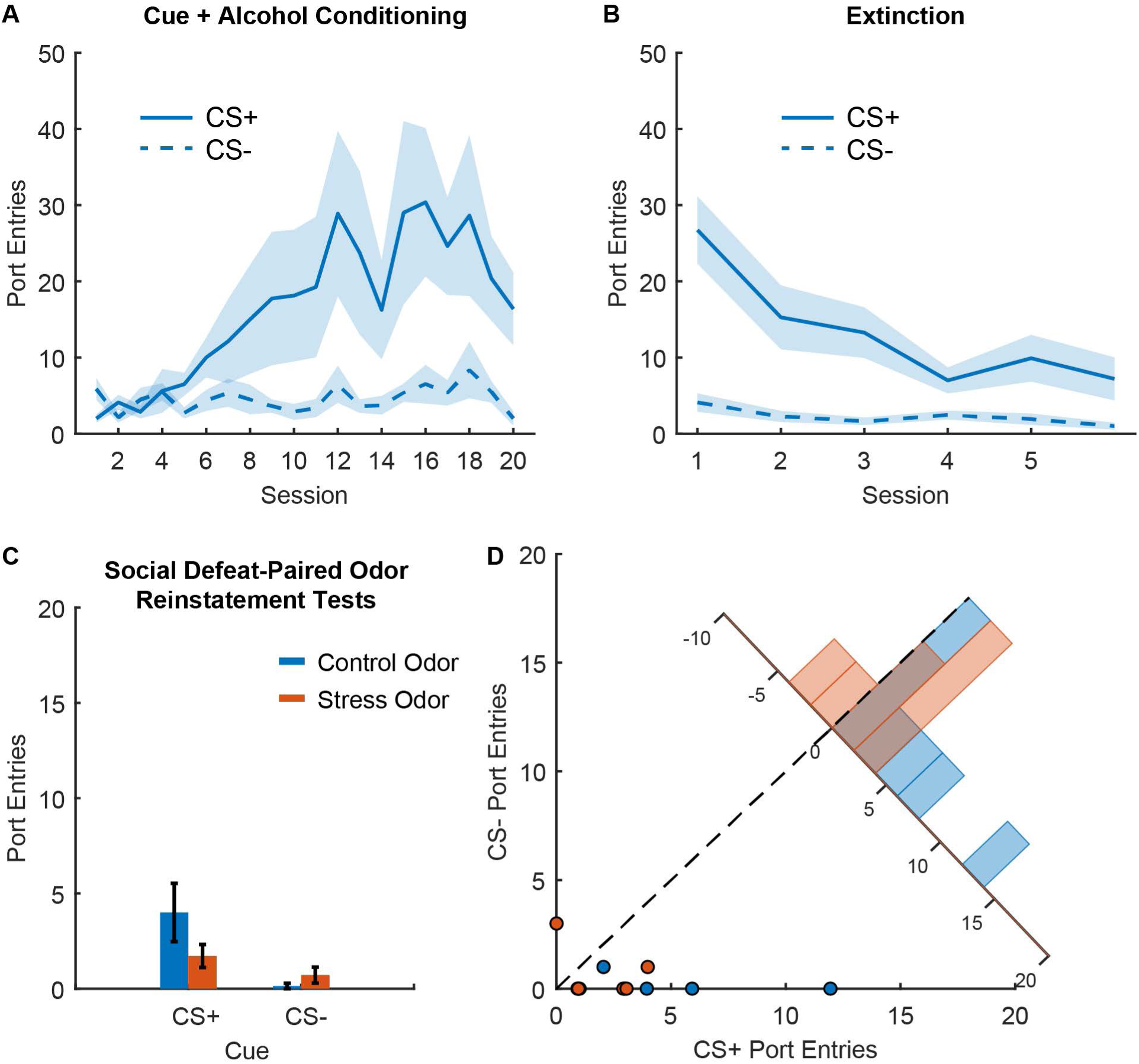
Reinstatement of responses to alcohol cues by olfactory cues associated with social defeat. A) Port entry number during the CS+ and CS-across Pavlovian conditioning with 20% ethanol reward (n=7), mean +/- SEM. B) Port entry number during the CS+ and CS-across extinction training, mean +/- SEM. C) Port entry number during reinstatement tests during the CS+ and CS-in presence of an olfactory cue previously paired with social defeat (stress odor, orange), or a control odor (blue), mean +/- SEM. D) Scatterplot showing port entry number during the CS+ and CS-from individual reinstatement sessions in presence of an olfactory cue previously paired with social defeat (stress odor, orange), or a control odor (blue). Inset histogram shows the distribution of sessions based on the difference in the number of port entries during the CS+ and the CS-.

Finally, we assessed reinstatement behavior in the presence of each rat’s social defeat odor cue or a control cue. In contrast to prior work with instrumental self-administration, we found that the social defeat stress odor failed to enhance Pavlovian conditioned responses to alcohol cues (Figure 5C and D). If anything, the odor paired with social defeat reduced port entries during the cue, though this effect did not reach statistical significance (main effect of stress odor, F(1,24) = 4.2066, p = 0.0513) which was not surprising given the low number of port entries in the control condition. We also found no effect of cue type (F(1,24) = 0.80516, p = 0.37847) or significant interaction between odor type and cue type (F(1,24) = 3.286, p = 0.082). We also saw a numerical reduction in the difference between port entry number during the CS+ and the CS-, but this effect also did not reach statistical significance (main effect of stress odor, F(1,12) = 2.9661, p = 0.11). We found no evidence of an effect on port entries during the intertrial interval (F(1,12) = 0.67, p = 0.43). Overall, the social defeat cues did not drive reinstatement of alcohol-seeking in the presence or absence of the Pavlovian cues and may have actually reduced behavior in this paradigm.

## Discussion

Here we found that while yohimbine can drive reinstatement of Pavlovian-conditioned responses, this effect does not generalize to other stressors that have been shown to reinstate instrumental alcohol seeking. Specifically, we found that yohimbine drove increases in port entry behavior after extinction of either cue-alcohol or cue-sucrose associations. While port entry behavior was greater during the CS+ during these tests, increases were also seen during the control cue and ITI periods. When we assessed the effects of intermittent footshock and odor cues paired with social defeat, we failed to see a reinstatement-like effect, and even saw a slight suppression of port entry behavior in the case of social defeat cues. Overall, our results suggest that the effects of stress on Pavlovian behavioral responses are complex and Pavlovian approach may not be the best model through which to understand the ways in which certain stressors potentiate cue-driven reward-seeking behavior.

### Yohimbine effects and their relationship to stress

One unanswered question from these studies is the relationship between the yohimbine-induced increases in alcohol seeking reported here and any stress-related effects of yohimbine. Yohimbine has been shown to elicit anxiety-like behaviors and increases in corticosterone in rodents (Davis et al. 1979; Johnston et al. 1988; Gill et al. 2013; Arrant et al. 2013), and yohimbine-induced reinstatement of instrumental alcohol and drug seeking behavior has been shown to depend on activation of extra-hypothalamic corticotropin-releasing factor receptors (Lê et al. 2000; Hansson et al. 2006; Marinelli et al. 2007b; Shalev et al. 2010). Despite this, the use of yohimbine as a stressor has been called into question in part due findings that suggest it is not aversive. Specifically, yohimbine does not increase the emission of 22 kHz vocalizations thought to indicate a negative affective state in rats (Mahler et al. 2013), and yohimbine produces a mild conditioned place preference (Chen et al. 2015). This critique conflates stress and aversion, and while stressors can certainly have negative affective qualities and long term negative affective consequences, stress is not defined by those negative affective qualities (Selye 1976). In some cases activation of stress systems may be rewarding; for instance, infusions of corticotropin releasing factor into the nucleus accumbens can also produce a conditioned place preference (Lemos et al. 2012).

An additional issue that has been raised for the use of yohimbine as a stressor is that it has been shown to cause reinstatement of operant responding for food, or even for cues alone (Ghitza et al. 2005; Nair et al. 2006; Chen et al. 2015). Yohimbine has also been shown to increase conditioned reinforcement by cues previously paired with sucrose or ethanol (Tabbara et al. 2020). While we did not replicate these conditioned reinforcement effects here, this could be due to differences in the length of Pavlovian training, or the fact that rats in our experiments did not experience the new operant-cue contingency prior to testing, which may have blunted overall responding across the testing conditions. Prior extinction may have also altered the ability of the cues to robustly reinforce novel operant responses. That being said, we did find similar effects of yohimbine on alcohol-and sucrose-seeking in our paradigm. This lack of specificity to drug-seeking behavior and drug-paired cues has contributed to uncertainty regarding yohimbine’s actions via stress-related mechanisms.

If stress generally acts as a context cue to drive reinstatement of drug-seeking (Schepers and Bouton 2019), it follows that stressors that reinstate drug-seeking should not reinstate food-seeking, since drug but not food reinforcers drive activation of stress systems (Sinha 2008). This is the case for intermittent footshock (Ahmed and Koob 1997; Buczek et al. 1999). A counterpoint to this is that food restriction, such as that implemented during Pavlovian conditioning with sucrose in this study, can also produce activation of stress systems (Stewart et al. 1988; Schroff et al. 2004), as does anticipation of food (Kalsbeek et al. 2012). Instead, the intensity of the stressor may also dictate whether that stress can act as an effective context cue for drug-seeking versus food-seeking. Stress may also act by activating dopamine systems and therefore increasing the motivational impact of reward-related cues (Robinson and Berridge 1993; Field and Powell 2007; Nair et al. 2011; Brown et al. 2012). If stress acts primarily via this mechanism to drive reinstatement, the question remains why many stressors fail to reinstate instrumental food-seeking behaviors, and why we failed to see reinstatement of Pavlovian approach after intermittent footshock or exposure to odors paired with social defeat.

### Stress intensity and null effects of intermittent footshock and social defeat cues

Reinstatement of reward-seeking by stress-induced activation of dopaminergic systems and resulting changes in the motivational impact of cues may depend on both the intensity of the stressor and the nature of the cue response being tested. This may explain in part why we failed to see an increase in Pavlovian approach after intermittent footshock and exposure to social defeat cues. Dopamine has been implicated preferentially in behavioral responses driven by the incentive value of cues, such as approach to a reward cue or “sign tracking,” in comparison to behaviors driven by the predictive value of cues, such as approach to a reward delivery location or “goal-tracking” (Flagel et al. 2011; Saunders and Robinson 2012). Here, we used auditory cues, which meant that the only measure of cue learning or motivated behavior we were able to monitor took the form of approach to the reward-delivery port, or goal tracking. Whether intermittent footshock or social defeat cues would potentiate or reinstate sign tracking behavior following Pavlovian conditioning remains unclear. Entry to a reward port where the reward is normally consumed may result in behavioral responses that are more closely tied to reward consumption, rather than reward-seeking. The effects of acute stress on alcohol consumption are complex (Becker et al. 2011). While it is difficult to identify specific rules that govern whether stress results in increases, decreases, or no change in alcohol consumption, more intense stressors may result in decreased consummatory behavior (Bond 1978; Champagne and Kirouac 1987; Van Erp and Miczek 2001; Darnaudéry et al. 2007), which could also affect goal-tracking behavior in response to cues. Therefore, even if intermittent footshock and social defeat cues can act to potentiate the motivational value of cues, these effects may be competing with opposing effects on consummatory behavior in our paradigm. It is also important to note that intense or prolonged stressors can also suppress even sign-tracking and reinstatement of instrumental drug-seeking behavior (Funk et al. 2005; Fitzpatrick et al. 2018).

## Conclusions

Here we found divergent effects of yohimbine, intermittent footshock and social defeat cues on reinstatement of behavioral responses to Pavlovian alcohol cues. Yohimbine potentiated alcohol seeking behaviors, whereas the other stressors failed to increase alcohol-seeking and may have reduced alcohol-seeking behavior. The results highlight the multiple behavioral mechanisms via which stress may act to increase or decrease drug-seeking behaviors. In some models of reward-seeking these stress effects and behaviors may be in conflict, which is distinct from the effects of a reward-paired context (Valyear et al. 2017). Pavlovian-conditioned approach may be a useful model for studying the excitatory effects of mild stressors, like yohimbine, on responses to drug-paired cues, but may be less useful to understanding that mechanisms by which more intense stressors potentiate the motivational value of drug-paired cues. Future work examining the effects of these stressors in other paradigms in which cues are presented non-contingently to drive incentive motivated behaviors is warranted.

## Acknowledgements

This work was supported in part by National Institutes of Health grants K99AA025384 and R00AA025384 to JMR, and a Provost’s Undergraduate Research Award to AA.

## Conflict of Interest

On behalf of all authors, the corresponding author states that there is no conflict of interest.

## Notes

### Competing Interest Statement

The authors have declared no competing interest.

